# *In vivo* fluorescence imaging with a flat, lensless microscope

**DOI:** 10.1101/2020.06.04.135236

**Authors:** Jesse K. Adams, Vivek Boominathan, Sibo Gao, Alex V. Rodriguez, Dong Yan, Caleb Kemere, Ashok Veeraraghavan, Jacob T. Robinson

**Affiliations:** Applied Physics Program, Rice University, 6100 Main Street, Houston, Texas 77005, USA; Department of Electrical and Computer Engineering, Rice University, 6100 Main Street, Houston, Texas 77005, USA; Department of Neuroscience, Baylor College of Medicine, One Baylor Plaza, Houston, Texas 77030, USA; Department of Bioengineering, Rice University, 6100 Main Street, Houston, Texas 77005, USA; Department of Computer Science, Rice University, 6100 Main Street, Houston, Texas 77005, USA

## Abstract

Fluorescence imaging over large areas of the brain in freely behaving animals would allow researchers to better understand the relationship between brain activity and behavior; however, traditional microscopes capable of high spatial resolution and large fields of view (FOVs) require large and heavy lenses that restrict animal movement. While lensless imaging has the potential to achieve both high spatial resolution and large FOV with a thin lightweight device, lensless imaging has yet to be achieved *in vivo* due to two principal challenges: (a) biological tissue typically has lower contrast than resolution targets, and (b) illumination and filtering must be integrated into this non-traditional device architecture. Here, we show that *in vivo* fluorescence imaging is possible with a thin lensless microscope by optimizing the phase mask and computational reconstruction algorithms, and integrating fiber optic illumination and thin-film color filters. The result is a flat, lensless imager that achieves better than 10 μm spatial resolution and a FOV that is 30× larger than other cellular resolution miniature microscopes.

## Introduction

The desire to understand how brain activity relates to animal and human behavior has driven the development of technologies that can capture neural activity with high spatial and temporal resolution over large areas of the brain. Fluorescence microscopy in particular shows great promise for measuring large-scale neural data with high resolution^1–4^. To apply fluorescence microscopy to freely moving animals; however, requires miniature microscopes with small lenses that can be mounted atop the head of a mouse (or other small animals)^5^. Miniature lens-based microscopes including two-photon microscopy^6^, widefield microscopy^7,8^, and optogenetic stimulation^9^ have all been demonstrated in the last few years. However, simultaneously achieving large FOV, cellular resolution (defined here as < 10 μm), and light weight (< 3g for freely moving mice) is extremely difficult. For example, previously reported miniature microscopes with cellular resolution have a FOV < 0.5 mm^2^ (Table 1)^5,6,10,11^. Miniature microscopes with a large FOV that can cover the entire mouse cortex must compromise spatial resolution to approximately 40 μm^8^. To reach the goal of simultaneously achieving a large FOV, and high resolution there are three potential approaches: (1) work with large animals that can support heavier weight^7^ (e.g. rat, monkey), (2) use external strain relief measures^12^, or (3) explore lensless designs (since lenses and housing account for a significant portion of device weight).

**Table 1.**
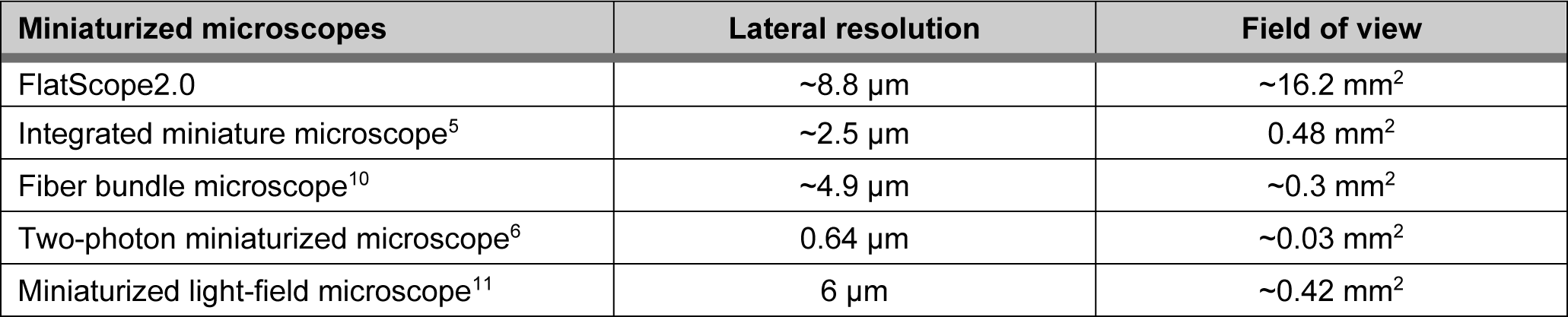
Comparison of FlatScope2.0 to miniaturized, head-mounted microscopes. Field of view and lateral resolution comparison of FlatScope2.0 to integrated, lens-based and fiber bundle microscopes. Selected microscopes have cellular resolution capability and remain light enough (< 3 g) to be mounted on a mouse and allow free behavior without any support.

We chose to overcome the physical limitations of lensed-based imaging systems by turning to computational imaging that uses joint design of optics, sensors and algorithms to open up new design degrees of freedom^13–15^. For example, computational imaging techniques have already made a significant impact on microscopy, for applications like super-resolution^16,17^, phase imaging^18^, and three-dimensional imaging^19,20^ that had previously been limited by constraints imposed by physical optics. Here we look to lensless microscopy, which completely replaces the traditional lens(es) of an optical system with computation^21^, or some combination of algorithms and light-modulating masks^22,23^. Unlike lens-based imaging systems where the goal is to project a reproduction of a (potentially magnified) scene onto an image sensor, a lensless imaging system seeks to produce an invertible transfer function between the incident light field and the sensor measurements. These measurements often may not resemble a traditional image^24–30^ but contain sufficient information for a computational algorithm to reconstruct an image. The major advantage for lensless imaging of fluorescence is the substantial benefits in FOV, three-dimensional capture, light-collection efficiency and form-factor^22–24,31^.

Despite the transformative potential of lensless microscopy, there are currently no demonstrations of *in vivo* fluorescence imaging due to two principle challenges: (a) reconstructing high-quality images in realistic, complex biological contexts and (b) integrating illumination and filtering in the reduces space between the sample and sensor. Previous lensless approaches have benefited from sparse, bright, or high-contrast samples where strong regularization and/or deblurring can be used to reconstruct estimates of the original image^22,31,32^. However, biological microscopy typically faces scenes that are dense, dim, and low contrast, which can often result in noisy reconstructed images. Additionally, previously lensless microscopes have primarily used trans-illumination to account for the fact that the sensor must be placed within a few millimeters of the sample to maintain high-resolution imaging; however, most *in vivo* applications require that the light source and sensor be on the same side in an epi-fluorescence configuration.

To address these two challenges, we designed FlatScope 2.0, which consists of a new, optimized point spread function (PSF) and light delivery system. We refer to this prototype as FlatScope2.0 because it is based on our previously demonstrated flat microscope^22^, but has significant design improvements that enables the first high-resolution *in vivo* fluorescence imaging using a lensless device.

The first major innovation in FlatScope 2.0 is a high-contrast and spatially sparse PSF that captures textural frequencies common in natural and biological samples (Fig 1a). To create this PSF we generate Perlin noise and apply uncanny edge detection^15^. Figure 1c shows the simulated modulation transfer function (MTF) of our contour-based PSF compared to other PSFs used in lensless imaging systems. The relatively flat MTF spectrum indicates that information from most spatial frequencies are well-preserved by the optical transfer function leading to improved image reconstruction for dense, low contrast samples like biological tissue. We chose to create this PSF using a phase mask (Fig. 1b), because they are capable of producing a wide variety of PSFs^33–38^ and allow higher light throughput compared to amplitude masks (which block some of the incoming light)^21^.

**Fig. 1.**
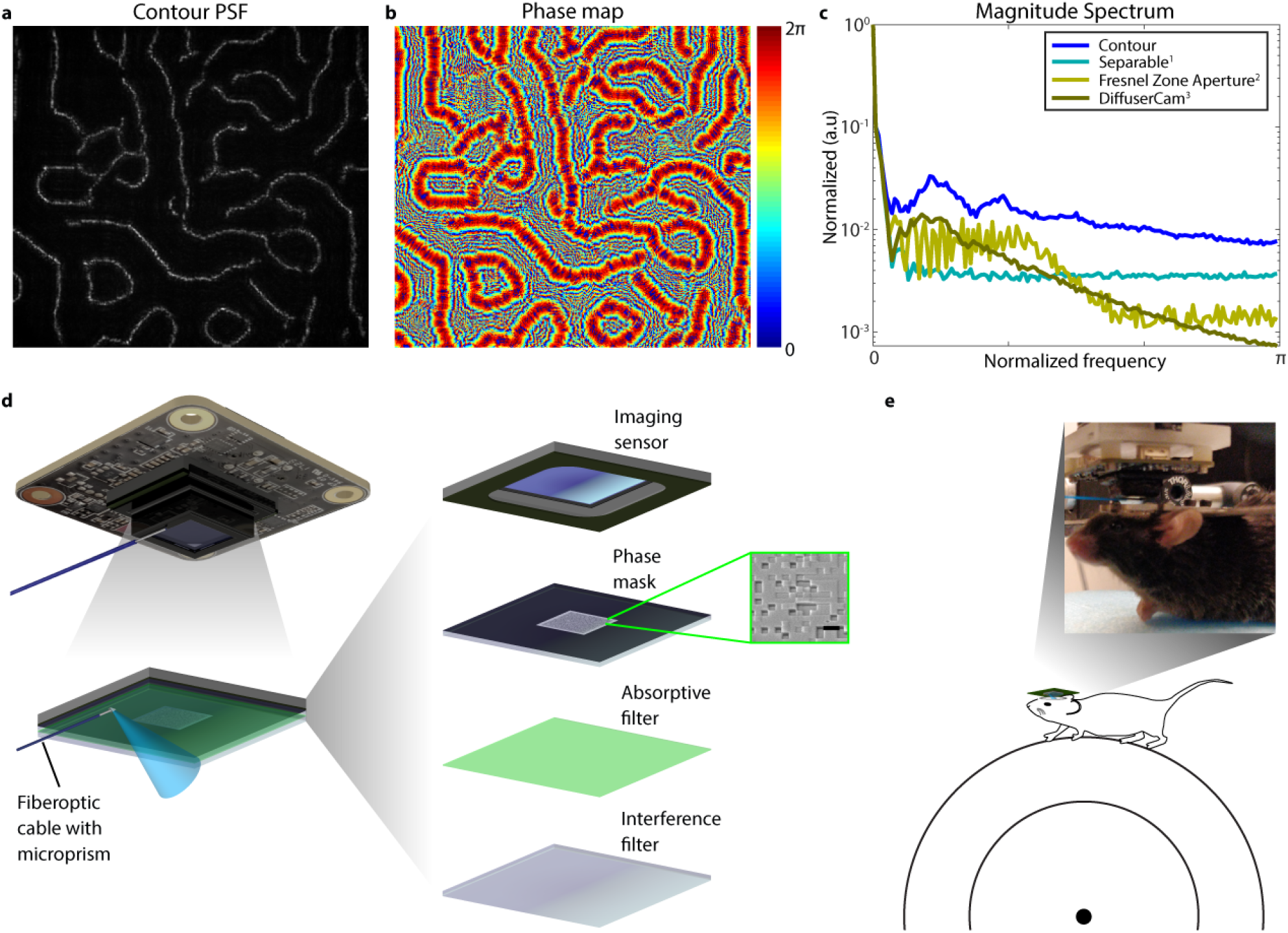
FlatScope 2.0 for *in vivo* fluorescence imaging. **a**, Contour-based PSF design provides robustness to noise and capturing many directional filters. **b**, Phase map for the mask in order to produce the PSF on the image sensor. **c**, Magnitude spectrum comparing PSF designs used by lensless imaging systems. **d**, FlatScope2.0 prototype including the off-the-shelf board level camera. Zoom-in shows the combined and exploded view of the components. Scanned electron micrograph of portion of phase mask. Scale bar 4 μm. **e**, Experimental setup for FlatScope2.0 for imaging a head-fixed mouse on a freely moving treadmill. Zoom-in shows a photo of a mouse on the treadmill with FlatScope2.0 in place.

The second major innovation in FlatScope 2.0 is a compact light delivery and filtering system that enables epifluorescence imaging. This system is based on a fiberoptic cable attached to an aluminum-coated microprism^39^. This illumination strategy is capable of fitting into the small space between the lensless imager and biological sample. To prevent the excitation light from reaching the sensor and overwhelming the fluorescence signal we constructed a hybrid excitation filter that combines an absorptive and interference filter^40^ and attached it to the top of the phase mask (see Methods). Figure 1d shows the prototype diagram with illumination method and component parts, the thickness between the image sensor and surface of device is ~5 mm.

## Results

To evaluate the performance of FlatScope2.0, we constructed prototypes using Sony IMX178 monochromatic 6 MP imaging sensors with 2.4-um pixels. Phase masks were fabricated using a two-photon lithography system (Nanoscribe, Photonic Professional GT2) for ease of prototyping (see Methods). The contour-based phase masks were designed with the minimum feature width of 12 μm.

### Spatial Resolution & Fixed Biological Samples

Using USAF resolution targets, we found that we could reconstruct images with < 9 μm resolution, which approaches the resolution needed to resolve individual neurons. We measured this resolution by capturing images of a negative 1951 USAF Resolution Target (Edmund Optics #59-204) with an added fluorescent background. Multiple images were captured at 15 ms each and averaged to remove noise. Excitation light was provided in a near-epi configuration with the fiber/microprism. Images were captured at a distance of ~4 mm. In Fig. 2a, we show that we can achieve < 9 μm resolution. While this spatial resolution is reduced by ~1.5× - 3.5× compared to (single-photon) miniature microscopes (see Supplementary Table 1), we find that our system achieves cellular resolution under sparse labeling conditions.

**Fig. 2.**
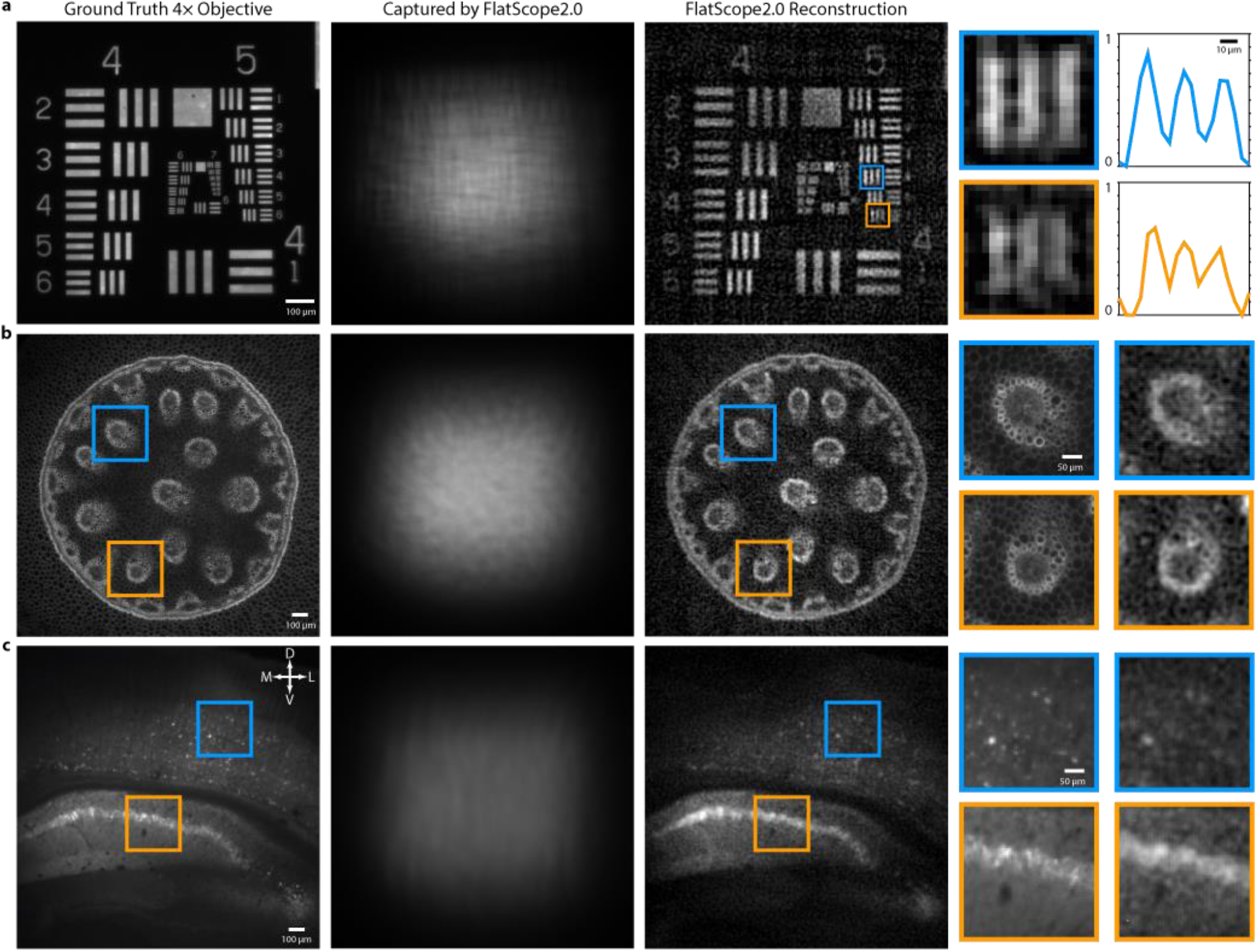
High spatial resolution in fixed biological samples. **a**, Ground truth capture by epifluorescence microscope, raw capture by FlatScope2.0, and FlatScope2.0 reconstruction of a USAF 1951 Resolution. Scale bar, 100 μm. Far right shows zoom-ins of group 5, elements of 4 & 6. Scale bar, 10 μm. **b**, Ground truth capture, raw capture by FlatScope2.0, and FlatScope2.0 reconstruction of a plant slice (*Convallaria majalis*). Scale bar, 100 μm. Far right shows zoom-in comparisons of ground truth and FlatScope2.0 reconstructions, respectively. Scale bar, 50 μm. **c**, Ground truth capture, raw capture by FlatScope2.0, and FlatScope2.0 reconstruction of a mouse brain slice expressing GCaMP6f. The compass shows dorsal-D, ventral-D, medial-M and later-L directions. Scale bar, 100 μm. Far right shows zoom-in comparisons of ground truth and FlatScope2.0 reconstructions, respectively. Scale bar, 50 μm.

We also found that the FlatScope2.0 can accurately reconstruct images of biological samples that are denser, dimmer, and lower contrast than resolution targets. FlatScope2.0 images of *Convallaria majalis* (lily of the valley) stained with green fluorescent protein (GFP) show good correspondence to ground truth images captured with a 4× microscope objective (Nikon Fluor) (Fig. 2b). This data is based on multiple images of the convallaria slice (at ~3.5 mm from the device) captured at 200 ms each in the near-epi configuration and averaged for noise removal. Although some detail is lost in FlatScope2.0 images compared to the ground truth images, we are still able to resolve some of the larger plant cells with sizes of around 10 μm. To further test FlatScope 2.0 with biological samples, we captured *ex vivo* images of a mouse brain slice (deep cortex and hippocampus) expressing the fluorescent protein GCaMP6f (see Methods). Here we capture 20 images at a distance of ~5.3 mm using transmissive illumination with 1 s exposures averaged to remove noise. Figure 2c shows a comparison of FlatScope2.0 to ground truth images. Similar to the image of convallaria, we see good reconstruction of the overall structure showing clear delineation between cortex layer 2/3, corpus callosum and hippocampus (CA1). We can also identify clusters of neurons in the cortex as well as the pyramidal layer of CA1 (see far-right column of Fig. 2c).

*In vivo* imaging presents additional challenges due to tissue scattering that causes blurring and loss of contrast. This blurring and loss of contrast affects the properties of the scene but has no direct effect on the performance of our reconstruction algorithm because the reconstruction algorithm makes no assumptions about the medium between the source and the imager. In fact, the blur and reduced contrast due to scattering are the same issues that affect epifluorescence imaging. The only additional effect on lensless imaging is the fact that noise is amplified by the reconstruction algorithm, which is true for all lensless imaging reconstruction techniques. The optimized FlatScope2.0 PSF with high contrast features provides good protection against this noise amplification and allows us to perform reconstructions through scattering media that are comparable with epifluorescence imaging (Supplemental Fig. 1).

### Single capture three-dimensional imaging *in vivo*

FlatScope2.0 is also capable of capturing three-dimensional information with a single image capture due to the fact that the PSFs change as a function of depth. To demonstrate this capability, we imaged Hydra vulgaris, a freshwater cnidarian polyp, which has a simple, yet dynamic network of neurons, optical transparency, and the ability to regenerate from a small patch of tissue^41,42^. An additional advantage of hydra is their genetic tractability and a number of transgenic lines including lines that express GFP or calcium dependent fluorescent protein GCaMP7b^43^. Using FlatScope2.0 we captured videos at 1 Hz of hydra expressing GFP in interstitial cell lineage. We were then able to reconstruct these images in three dimensions during post-processing using a series of PSFs corresponding to different distances from the sensor in 50 μm increments (Fig 3a). Here we show a 4 mm^3^ volume that we reconstructed from a single capture. By contrast traditional microscopes typically require scanning spatially, axially, or both. This single-capture, three-dimensional imaging opens up the possibility for high-resolution, high-speed, volumetric imaging of whole organisms freely behaving over large FOVs.

**Fig. 3.**
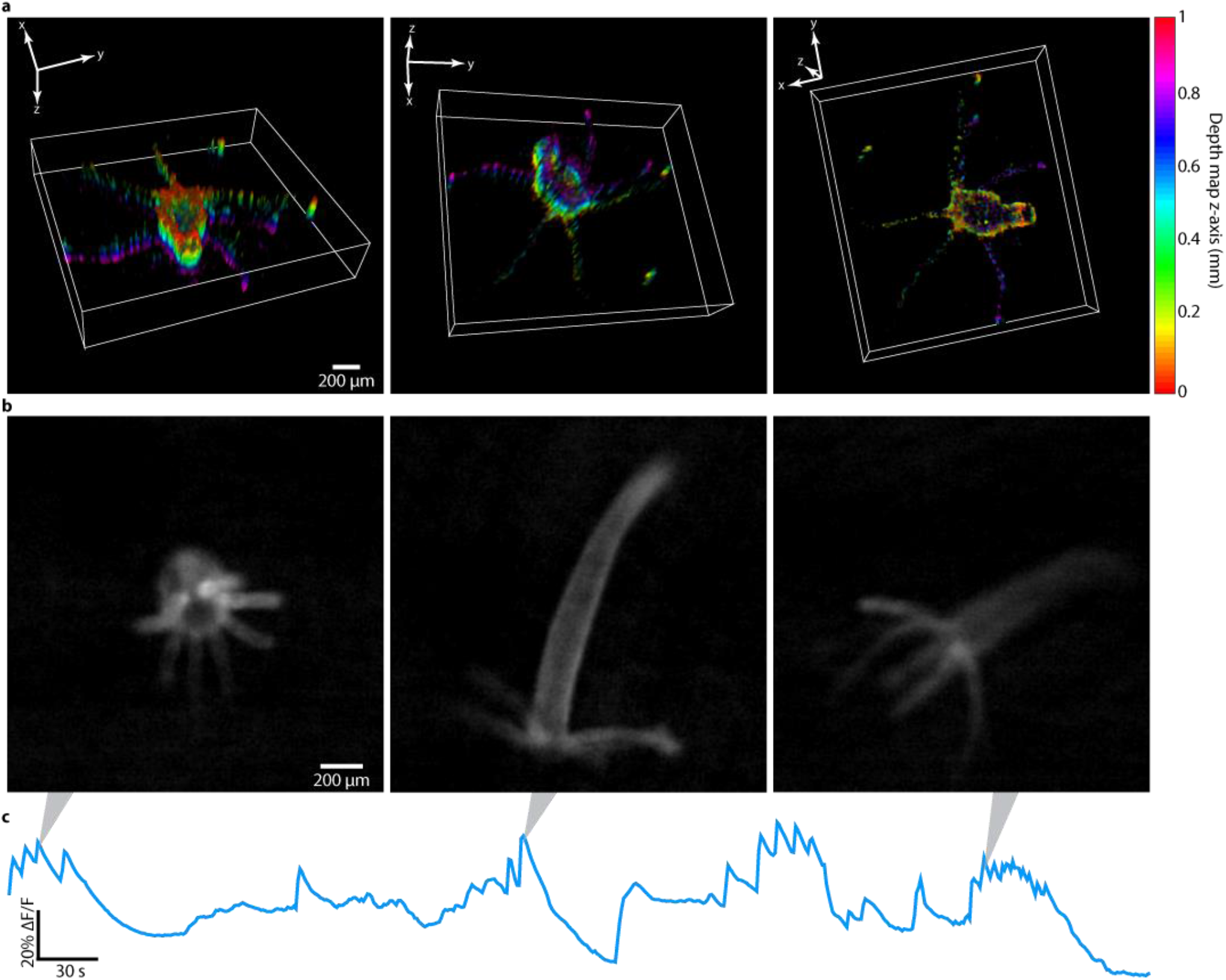
Ca^2+^ imaging (GCaMP7b) and three-dimensional imaging (GFP) of Hydra vulgaris *in vivo*. **a**, Select angles of three-dimensional FlatScope2.0 reconstructions of Hydra vulgaris expressing GFP in the interstitial cell lineage. Colors represent relative depth location along the z-axis. Scale bar, 200 μm. **b**, Selected FlatScope2.0 reconstructed frames of video of Hydra vulgaris expressing GCaMP7b in muscle cells. Scale bar, 200 μm. **c**, ΔF/F traces showing the Ca^2+^ responses over a 5-minute recording.

### *In vivo* epifluorescence calcium imaging of whole animal with *Hydra vulgaris*

In addition to reconstructing structural images based on GFP, we were able to reconstruct *in vivo* calcium dynamics by imaging genetically encoded calcium indicators (GECIs). We began by imaging *Hydra vulgaris* expressing GCaMP7b in muscle cells. We captured video of hydra freely behaving at 2 Hz over a FOV > 12 mm^2^. In Fig. 3b&c we show selected frames from a video of the reconstruction as well as the overall change in fluorescence over time (ΔF/F). The recordings show strong calcium responses during contraction events as reported previously^44^ (full video shown in Supplementary Video 1). Although movement of the animal can produce some noise in the ΔF/F signal, the deformable *Hydra* body makes it difficult to generate adaptive ROIs that move with the animal. Nevertheless, the bright synchronous calcium activity in the Hydra peduncle allows for fixed ROIs to effectively capture calcium dynamics^41,45^. The videos reconstructed using FlatScope2.0 can be used for studying the behavior of microorganisms and the associated calcium responses. Additionally, the FOV captured shows the potential for ultra-widefield imaging using this lensless system approach.

### *In vivo* epifluorescence calcium imaging in mice

We also found that FlatScope2.0 can capture calcium dynamics from mouse cortex that is comparable the dynamics recorded using epifluorescence microscopy. Mice expressing GCaMP6f in the motor cortex (see Methods) were head-fixed and placed upon a freely moving treadmill to allow for locomotion during video capture and velocity data was recorded (synchronized with frame captures). We recorded calcium imaging movies over a FOV of ~16 mm^2^ of the entire cranial window (Supplementary Fig. 2) using a 473 nm excitation light source (see Methods). A cropped region near the injection site is shown in Figure 4a. Compared to previously reported miniature microscopes with cellular resolution, we achieve more than a 30× increase in the FOV (Table 1). During the recording sessions, we applied a brushing tactile stimulation in 30 s intervals to encourage locomotion and neural response in the motor cortex. Correlation between locomotion (as performed by the mouse here on a treadmill) and neuronal response in the motor cortex have been established in prior reports^46–48^ and are confirmed in our epifluorescence imaging. In the regions of high activity in the motor cortex, we observed peak fluorescence signals during locomotion an average of ~3× greater than periods of no motion (rms) and were able to resolve blood vessels as small as ~10 μm in diameter. To compare our FlatScope2.0 imaging to conventional epifluorescence, we replaced the FlatScope2.0 with a 4× objective lens and fluorescence microscope. We performed this epifluorescence imaging under the same conditions within 30 minutes of our FlatScope2.0 recordings (see Methods) and found very similar spatiotemporal dynamics in the two data sets (Fig. 4 and Supplementary Fig. 3 & 4). We chose the 4× objective lens to accommodate the FOV captured by FlatScope2.0 (while maintaining high resolution) and registered both data sets to accommodate for movement^49,50^.

**Fig. 4.**
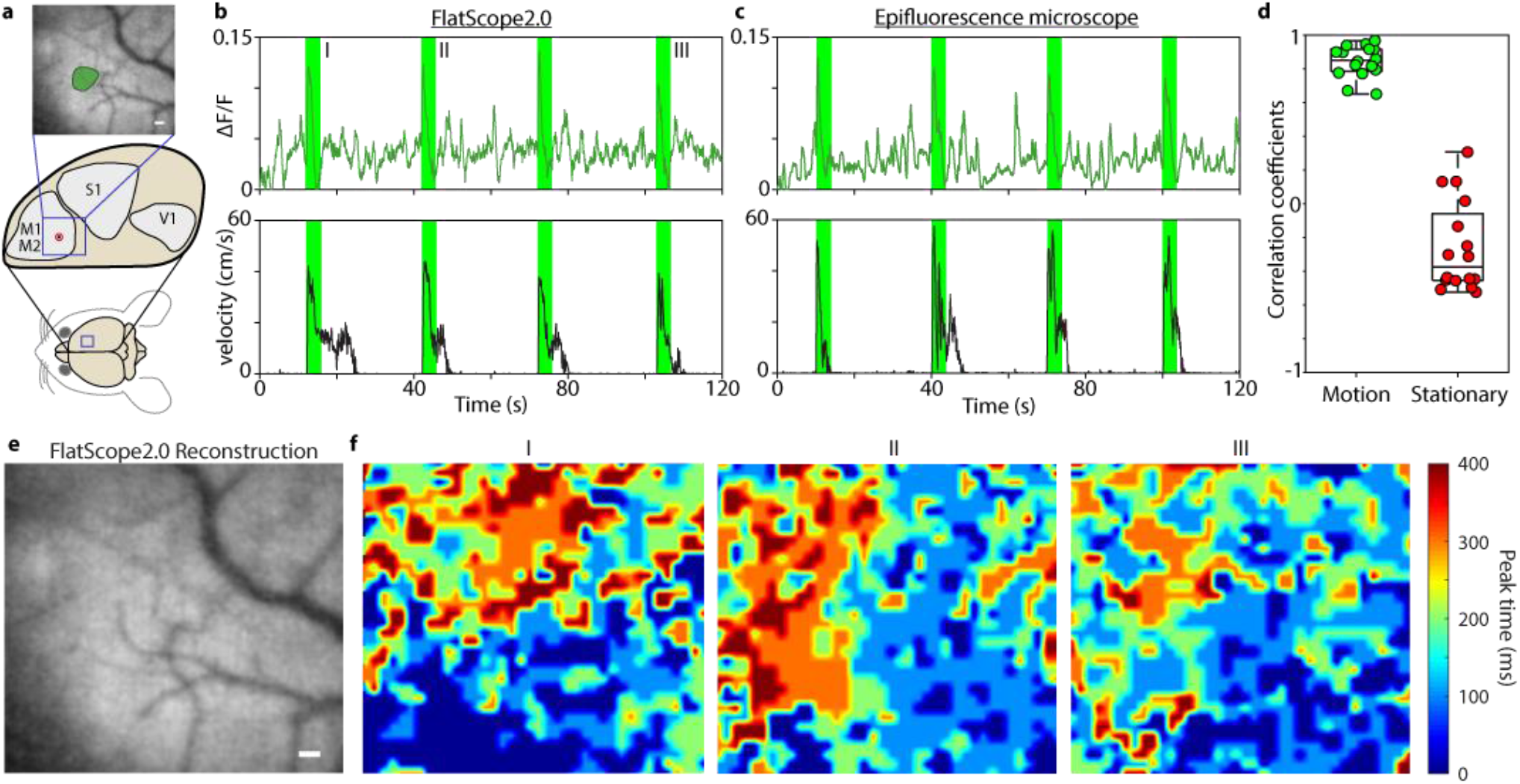
Comparison of FlatScope2.0 to epifluorescence during stimulus-evoked Ca^2+^ responses in motor cortex. **a**, Target region in the mouse brain. The blue square region indicates the approximate location of the cropped FOV (in the motor cortex). The circle with the black dot indicates the approximate region of injection for GCaMP6f. Zoom-in shows a FlatScope2.0 reconstruction with a single ROI of high-activity marked. Scale bar, 100 μm. **b**, ΔF/F trace and treadmill velocity for FlatScope2.0 during recording session. The rising edge of the 4-second windows in green correspond to the application of tactile stimuli. **c**, ΔF/F trace and treadmill velocity for epifluorescence during recording session. **d**, Box plot showing correlation coefficients comparing stimulus-evoked Ca^2+^ responses from FlatScope2.0 to epifluorescence (shown in green), and comparison of periods of little or no motion with FlatScope2.0 to stimulus-evoked Ca^2+^ responses with epifluorescence (shown in red) for the ROI. **e**, FlatScope2.0 reconstruction. Scale bar, 100 μm. **f**, Heat maps for FlatScope2.0 reconstructions showing spatiotemporal Ca^2+^ dynamics time-aligned with stimuli (at I, II, and III). Colormap shows the time at which pixels have their peak response for ΔF/F during a 400 ms period.

To compare lensless with epifluorescence data, we selected 4-second windows time-aligned to the response to stimulation (reflected as a large change in ambulatory activity of the mouse) over a two-minute period (Fig. 4b & c, shown in green). Ambulatory activity was determined by synchronizing the wheel velocity with the imaging data for both FlatScope2.0 and epifluorescence (shown in Fig. 4b & c for a single ROI, Supplementary Video 2 shows multiple regions). We validated the similarity of the windowed ΔF/F responses with correlation, finding a median correlation coefficient of 0.852 with interquartile range (IQR) of 0.131 (Fig 4d, left-side). We also compared data from time windows of little to no locomotion, where velocity < 1 cm/s (windows shown in Supplementary Fig. 3) with epifluorescence captured motion resulting in a median correlation coefficient of −0.375 and an IQR of 0.513 (Fig 4d, right-side). As expected, data from regions of motion/stimuli highly correlate when comparing FlatScope2.0 to epifluorescence data (coefficients range from 0.65 to 0.97, *p* < 0.001), while regions of little to no motion show more random correlation (coefficients ranging from −0.52 to 0.31, two values of *p* < 0.001 and six values of *p* > 0.05), confirming that using FlatScope2.0 we can capture behaviorally relevant calcium dynamics that are comparable to traditional dynamics capture with conventional epifluorescence microscopes.

In addition, when plotting the time of the calcium peaks over the FOV we find that when the animal initiates movement the majority of early peaks appear in the bottom and right of the FOV and the majority of late peaks appear in the top and left (Fig. 4f). This pattern is also observed with epifluorescence imaging confirming that FlatScope measures similar spatiotemporal Ca^2+^ dynamics (images are processed for visualization, see Supplementary Fig. 4 for epifluorescence data). The recorded video was captured at 10 Hz (50 ms exposures) and shows a period of 400 ms following the tactile stimulation.

We can also extract unique calcium activity from small regions of interest (ROIs) that approach the size of individual cells. To capture this data, we rely on the fact in the time domain, some Ca^2+^ activity in individual or small groups of cells appear as outliers compared to the background activity. Robust Principle Component Analysis (RPCA) excels at determining such sparse outliers^51,52^, and so we used the RPCA algorithm to identify these regions of high calcium activity. We then used K-means to identify which high-activity regions show correlated activity. These clustered ROIs represent distinct calcium signals from areas that are the size of a few individual cells. When we compared the ROIs determined through RPCA and k-means from FlatScope2.0 we found close correspondence to bright individual neurons observed from epifluorescence microscopy (Fig. 5d). Additionally, the calcium activity in each of these ROIs show unique temporal dynamics suggesting that calcium dynamics might be recovered with near cellular resolution.

**Fig. 5.**
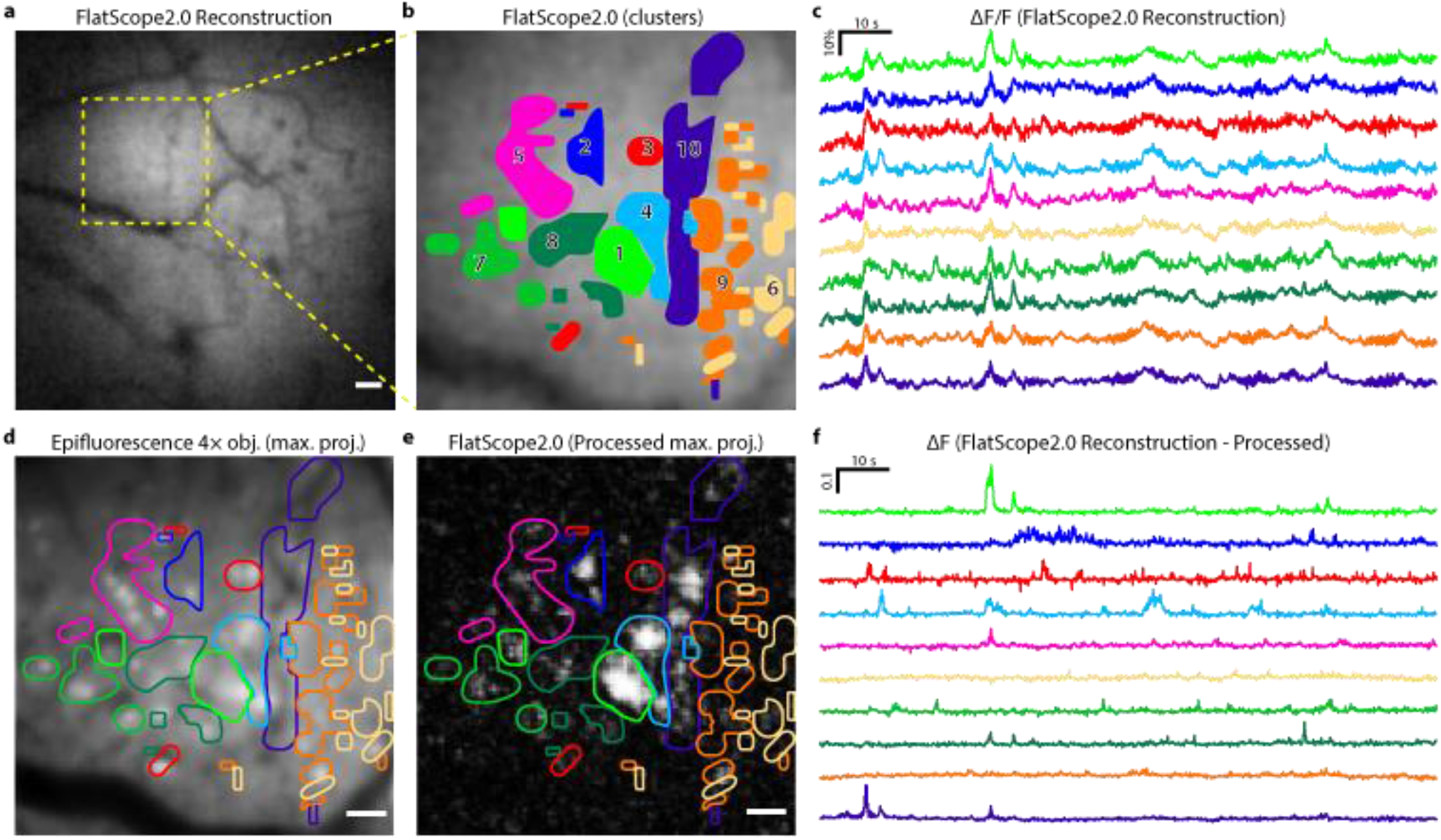
Extracting cell-sized Ca^2+^ signals from FlatScope2.0 reconstructions. **a**, Cropped region of FlatScope2.0 reconstruction of a single frame with a high-activity region shown by a dashed box. Scale bar, 100 μm. **b**, Zoom-in on the region of high-activity with overlay of clusters determined through post-processing using RPCA and k-means. Scale bar, 50 μm. **c**, ΔF/F traces from FlatScope2.0 reconstruction video over during two-minute recording corresponding to the clusters. **d**, Maximum projection from epifluorescence recording of same high activity region with overlay of clusters determined from FlatScope2.0 data. Scale bar 50 μm. **e**, Maximum intensity projection of FlatScope2.0 reconstruction data processed using RPCA with overlay of clusters. Scale bar 50 μm. **f**, ΔF traces from FlatScope2.0 reconstructions after post-processing using RPCA.

## Discussion

We have demonstrated structural and functional fluorescence imaging of biological samples using lensless microscopes with cellular resolution (reaching < 9 μm with a resolution target), FOVs >16 mm^2^, digital refocusing, and single capture, three-dimensional imaging over volumes of 4 mm^3^. We have also shown, to the best of our knowledge, the first *in vivo* Ca^2+^ imaging with a lensless microscope. From this calcium imaging we can extract localized Ca^2+^ signals ranging in area from 150 μm^2^ to 0.01 mm^2^ using RPCA. These results serve as a proof-of-principle that lensless microscopy provides a route toward high-resolution and large FOV calcium imaging in a small lightweight form factor. By reducing (or rearranging) the electronic components around the imaging sensor, we foresee miniaturizing our device to the point that we can mount it atop a mouse or small animal to study calcium activity during unrestricted behavior.

While the resolution shown here is at the level of individual cells, improvements are needed to increase the resolution to the sub-cellular level. This improvement could be realized by optimizing the masks for closer scene-to-mask distances. These closer distances will not only improve spatial and axial resolution but will increase light collection efficiency and reduce the form-factor enabling easier integration. Additionally, integration of the device can allow for a smaller refractive index change between the sample and device surface, which can cause a slight barrel effect at the edge of the FOV. As one brings the device closer to the sample, there is a corresponding decrease in the FOV until it matches the size of the sensor. To increase the effective FOV, future devices could be arrayed to maintain large scene coverage while maintaining a thin device profile.

Other modifications to the design could further improve performance. Integration of the excitation light source (or multiple sources) can be introduced to improve the illumination coverage over the FOV. For this purpose, commercially available micro-LEDs or integrated waveguides can be used at little to no cost to form factor. To improve on the low-light capabilities necessary for quality fluorescence microscopy, sCMOS imaging sensors or single photon avalanche diodes (SPADs) could be integrated to increase the SNR which is so important in lensless imaging. In addition to physical alterations of the lensless system, integration of new computational methods for imaging through scattering media might be used to extract additional three-dimensional information for brain imaging. One important opportunity for this class of lensless *in vivo* imagers is the potential to image tens of thousands of neurons simultaneously in a freely moving animal. Additionally, the principles behind FlatScope2.0 do not limit our imaging to fluorescence microscopy. Because our system is designed to capture incoherent sources, other imaging modalities like brightfield, darkfield and reflected-light microscopy are also possible, which could be useful for applications including endoscopy and point-of-care diagnostics. Overall, technologies based on FlatScope2.0 can be applied to a number of challenging microscopy problems where high temporal resolution, large FOV, and small form-factor are outside the capabilities of traditional lens-based microscopy.

## Methods

### Calibration

The PSF of the mask must be learned prior to capturing scenes through a one-time calibration process. Because we cannot capture images of an ideal point source, we instead use a single 10um fluorescent microsphere (FluoSpheres yellow-green), which is closely aligned to the center of our mask. The fluorescent bead was dropcast onto a microscope slide, then protected by a coverslip (Nexterion, 170 μm thick). Images of the PSF were captured for each depth (i.e. distance axially from the device) of interest by FlatScope2.0 with the bead located approximately at the center of the phase mask. Calibration images were averaged through multiple captures and background was subtracted to ensure the highest SNR for the PSFs. The sensitivity of the system to depth requires calibration images of PSFs to be taken over a range of distances, which allow for refocusing and three-dimensional reconstruction in post-processing.

### Reconstruction

To reconstruct images, we effectively solve the following minimization problem, adding Tikhonov regularization to the deconvolution to avoid noise amplification:

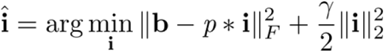

where * denotes convolution, 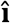 is the estimate of the scene, **i** is the scene, **b** is the sensor measurement, *p* is the PSF at the scene distance, *γ* is the regularization weight, and F is the Frobenius norm. This minimization problem can be solved by closed form via Wiener deconvolution as:

where ⨀ denotes Hadamard product, 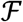 denotes a Fourier transform and 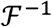 denotes an inverse Fourier

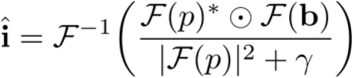

transform. To reconstruct three-dimensionally, we solve the same minimization problem, but do so for the sum of distances of interest:

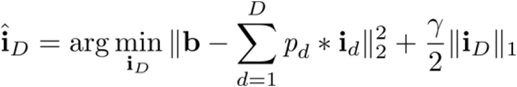

where *D* is the number of distances, *d*, from the imaging device.

### Excitation Light

Excitation light for FlatScope2.0 was achieved both transmissively and in near-epi configurations. Transmissive illumination was provided either using light through an epi-fluorescent microscope objective (Nikon Fluor 4×, NA 0.13 PhL DL), or using a near-collimated 470 nm LED (Thorlabs, M470L3) with an incorporated GFP excitation filter (Thorlabs, MF469-35). Near-epi illumination was provided using a fiber-coupled 475 nm LED (Prizmatix, UHP-T-475-SR) with an incorporated GFP excitation filter (Thorlabs, MF469-35). The light was coupled into a multimode fiberoptic cable (Thorlabs M72L01 200um diameter core, NA 0.39 or Edmund Optics, #57-749 400 μm diameter core NA 0.22) with an aluminum coated microprism (Edmund Optics 0.18 mm #66-768 or 0.70 mm #66-773) adhered to the exposed tip of the fiber with optical epoxy (Norland, NOA72). The microprism was placed near the surface of the lensless imaging device (Fig. 1d).

### Fabrication

FlatScope prototypes were constructed using an off-the-shelf board level camera (Imaging Source, DMM 37UX178-ML) with a monochrome Sony CMOS imaging sensor (IMX178LLJ, 6.3 MP, 2.4 μm pixels). The phase mask was fabricated using a 3-D photolithographic system (Nanoscribe, Photonic Professional GT2) using a photoresist (Nanoscribe, IP-L 780) on a 700 μm SiO2 substrate. The substrate was then cut down to more closely match the sensor size of the camera (and filters). The substrate was masked with an opaque material to provide an aperture containing only the phase elements. The substrate with phase mask was affixed atop the imaging sensor, followed by a hybrid filter, similar to Richard et al.^40^, but using a commercially available absorptive filter (Kodak, Wratten #12) placed below an interference filter (Chroma, ET525/50m), both cut to the sensor size requirements (see Fig. 1d). A housing was 3D printed (MJP 2500) to hold the phase mask and filters atop the imaging sensor.

### Mouse Brain Slice

All experiments were approved by the Rice University IACUC. Mice were sacrificed 14 days post-injection of AAV9-CamKII-GCaMP6f in the CA1 region of the hippocampus. From bregma, the injection site was +2 mm medial/lateral, −2 mm anterior/posterior and −1.65 mm deep. The total injection amount was 0.5 μL at a rate of 0.05 μL/min. The brain expressing GCaMP6f was cryoprotected in 30% sucrose, embedded in OCT compound (Tissue-Tek), frozen and sliced into 50 μm slices using a cryostat (Leica). Slices were then rinsed in 1x phosphate-buffered saline (PBS) and mounted using Vectashield H-1000 with DAPI (Vector Labs) then sealed with coverslips.

### Hydra vulgaris

Hydra were cultured in Hydra media using the protocol adapted from Steele Lab (UC Irvine) at 18°C with 12:12 hours light-dark cycle. Animals were fed freshly hatched artemia every 2 days and cleaned after 4 hours. Transgenic lines developed by embryo microinjections^43^ expressing either GFP in interstitial cell lineage (GFP, Neurons) or GCaMP7b in endodermal muscle cells (GCaMP7b, Endo) were used. The transgenic line of Hydra expressing GFP under actin promoter was originally developed by Steele Lab and selected for expression in neurons. The transgenic line of Hydra expressing GCaMP7b under EF1a promoter was developed by Robinson Lab and Juliano lab (UC Davis) selected for expression in endodermal muscle cells.

Both hydra expressing GFP and GCaMP7b were imaged in hydra media on petri dishes with coverslip bottoms (MatTek, 35 mm, 14 mm #1.5, coverslip). FlatScope2.0 images were captured through the coverslip with excitation light provided by the fiberoptic cable/microprism combination.

#### Brain tissue phantom

The brain tissue phantom was created by suspending non-fluorescent polystyrene microspheres (Polybead Acrylate Microspheres, 1 μm diameter, 4.55 × 10^10^ beads/mL) in polydimethylsiloxane (PDMS) (Sylgard, Dow Corning; 10:1 elastomer:cross-linker weight ratio). For the brain tissue phantom, a concentration of 5.46 × 10^9^ beads/mL was used to achieve a scattering coefficient of ~1 mm^−1^ to simulate the reported scattering coefficients in mouse brain tissue^53,54^. Microspheres (0.6 mL) were added to isopropyl alcohol (0.6 mL) and vortexed thoroughly for uniformity. The microsphere/alcohol mixture was added to 4 g of the PDMS elastomer and mixed thoroughly, followed by 0.4 g of crosslinker and again mixed thoroughly for uniformity. The desired thickness of 140 μm was achieved by spin-coating the mixture onto a SiO2 wafer. then heated at 37 °C for 30 minutes.

### *In vivo* mouse brain imaging

#### Animals

Wild-type C57BL/6 mice (n=3) from Charles River Laboratories were used for this study. Animals are housed with standard 12 h light/dark cycle with ad libitum food and water. All animals (n=3) were injected with the adeno-associated viral vector AAV9.CamKII.GCaMP6f.WPRE.SV40 (Penn Vector Core). All experimental procedures were approved by the Institutional Animal Care and Use Committee at Rice University and followed the guidelines of the National Institute of Health.

#### Headpost Implant and Cranial Window Design

Headpost implant design was adapted from Ghanbari et al.^55^, and consists of a custom-made Titanium or Stainless-steel head-plate, a 3D-printed frame, and three 0-80 screws to hold the frame to the head-plate. We fabricated the head-plate with Titanium or Stainless-steel plate (McMaster-Carr) using a waterjet system (OMax), and 3D printed the frame using a ProJet MJP 2500 (3D Systems). Our design files can be found online (https://github.com/ckemere/TreadmillTracker/tree/master/UMinnHeadposts). We assembled the headpost implant after tapping the 3D printed frame with 0-80 tap and securing the head-plate over the frame with three screws. The entire headpost is then stored in 70% Ethanol prior to surgery.

Cranial window fabrication procedure was adapted from Goldey et al.^56^. Windows were made of 2 stacked round coverslips (Warner Instruments # CS-3R, CS-4R, CS-5R) of different diameters. To fabricate the stacked windows, a 3 mm (or 4 mm) round coverslip was epoxied to a 4 mm (or 5 mm) cover slip using an optical adhesive (Norland Products Inc. e.g. # NOA 61, 71, 84) and cured using long-wavelength UV light. To accommodate the large 5 mm stacked window, we cut off the right side of the 3D-printed frame to allow for extra space for the C&B Metabond to bind to the skull outside of the stacked cranial window. Fabricated stacked windows were stored in 70% ethanol prior to surgery.

#### Surgical Procedures

For AAV9 injections, mice were anesthetized with 1-2% isoflurane gas in oxygen and administered sustained release Buprenorphine SR-LAB (Zoofarm, 0.5 mg/kg). A small craniotomy was carefully drilled at the target location. A total of 0.5-1 μL of AAV9 virus was injected slowly at a rate of 0.07 μL/min into each mouse with a Hamilton syringe paired with syringe pump controller (KD Scientific # 78-0311). Specifically, mouse W1 was injected at −1.5 AP; +1.5 ML; −0.25 DV targeting the motor cortex; mouse W2 was injected at −1.67 AP; +1.1 ML; −0.25 DV targeting the motor cortex; mouse W3 was injected at −1.23 AP; +1.23 ML; −0.25 DV targeting the motor cortex. Following the injection, a small amount of bone wax was applied over the craniotomy, while taking care not to press down on the brain surface. The incisions were closed using a small drop of Vetbond (3M), and the mice were allowed to express for at least 4 weeks before headpost and window implantation.

For headpost and window implantation, mice were administered Buprenorphine SR-LAB (0.5 mg/kg) and Dexamethasone (2 mg/kg) 30 min prior to the craniotomy procedure. Mice were anesthetized with 1-2.5% of isoflurane gas in oxygen and secured on a stereotax (Kopf Instruments) using ear bars. A single large cut was made to cut away the majority of the skin above the skull while attempting to match the 3D printed frame to the exposed skull. We then used a 3 mm (or 4 mm, depending on the window implant size) biopsy punch to carefully center the craniotomy over the AAV injection site while avoiding the sagittal suture, and slowly and gently rotated the biopsy punch until the bone could be lifted away with 5/45 forceps. We used saline to irrigate regularly while performing the craniotomy, taking care not to puncture the dura. After stopping any bleeding, we placed a small amount of silicone oil (Sigma-Aldrich Inc. # 181838) to cover the brain, then carefully placed the stacked window over the brain. We then applied pressure to the top of the stacked window with a thinned wooden applicator (e.g. back of a cotton swab or toothpick) mounted on the stereotaxic arm until the 4 mm (or 5 mm) coverslip was flush with the skull surface. We used cyanoacrylate to glue around the window and waited until it dried before removing pressure from the thinned wooden applicator. We then positioned the headpost over the skull and used C&B Metabond to cement the headpost in place, taking care not to cover the cranial window. After the Metabond dried, we applied some silicone elastomer (World Precision Instruments, Kwik-Sil) over the cranial window to protect it from any damage. We then administered post-operative drugs (meloxicam at 5 mg/kg and 0.25% Bupivicaine around the headpost implant) and allowed the mouse to recover for at least 3 days before imaging.

#### Recording sessions

Prior to each animal’s first awake head-fixed imaging session, the animals were acclimated to head-fixation and the treadmill set-up for at least three sessions. Additionally, to minimize stress prior to awake imaging session, we acclimated the animals to mechanical fixation in which the head was restrained. Animals received chocolate milk (Nesquik) as a reward while running on a custom-built treadmill. Once acclimated, animals were imaged for up to two sessions per day, where each session lasted no longer than two hours.

For experiments, mice were head-fixed atop a freely moving treadmill during video capture. Epifluorescence microscope images were captured with a sCMOS camera (Kiralux, 5 MP) through a 4× objective. Treadmill rotation data (synchronized with the camera) was captured during the experiment. During the recording session, a brushing tactile stimulus was applied to the spine region of the mouse with a minimum of 30 s between stimulus events. FlatScope2.0 recording sessions were captured within 30 minutes of epifluorescence recordings, with exposure time, recording lengths and stimuli timing matching those with the epifluorescence microscope. ROIs in the brain for comparing treadmill activity across epifluorescence and FlatScope2.0 were selected based on high activity areas extracted from the epifluorescence video.

#### RPCA analysis

This analysis was based on two-minute imaging sessions recorded at 20 Hz over the full FOV of the cranial window using both FlatScope2.0 and an epifluorescence microscope with a 4× objective. The ROIs for cluster analysis were selected by observing a 500 μm x 500 μm region of spontaneous high activity in the FlatScope2.0 reconstructions (Fig. 5a). This same ROI was used for both FlatScope2.0 and epifluorescence data. Figures 5b & c show the clusters extracted through k-means as well as the corresponding ΔF/F for those clusters.

## Acknowledgements

We would like to thank Krishna Badhiwala for the useful discussions on *Hydra* neurobiology and behavior. We would also like to thank Jill Juneau for the mounted mouse brain slices

## References

1. Prevedel, R. et al. Fast volumetric calcium imaging across multiple cortical layers using sculpted light. Nature Methods vol. 13 (2016).

2. Sofroniew, N. J., Flickinger, D., King, J. & Svoboda, K. A large field of view two-photon mesoscope with subcellular resolution for in vivo imaging. Elife 5, (2016).

3. Dombeck, D. A., Khabbaz, A. N., Collman, F., Adelman, T. L. & Tank, D. W. Imaging large-scale neural activity with cellular resolution in awake, mobile mice. Neuron 56, 43–57 (2007).

4. Tomer, R. et al. SPED Light Sheet Microscopy: Fast Mapping of Biological System Structure and Function. Cell 163, 1796–1806 (2015).

5. Ghosh, K. K. et al. Miniaturized integration of a fluorescence microscope. Nat Methods 8, 871–878 (2011).

6. Zong, W. et al. Fast high-resolution miniature two-photon microscopy for brain imaging in freely behaving mice. Nat Methods 14, 713–719 (2017).

7. Scott, B. B. et al. Imaging Cortical Dynamics in GCaMP Transgenic Rats with a Head-Mounted Widefield Macroscope. Neuron 100, 1045—-1058.e5 (2018).

8. Rynes, M. et al. Miniaturized device for whole cortex mesoscale imaging in freely behaving mice. in Neural Imaging and Sensing 2020 (eds. Luo, Q., Ding, J. & Fu, L.) vol. 11226 15 (SPIE, 2020).

9. de Groot, A. et al. Ninscope, a versatile miniscope for multi-region circuit investigations. eLife 9, (2020).

10. Kim, M., Hong, J., Kim, J. & Shin, H. Fiber bundle-based integrated platform for wide-field fluorescence imaging and patterned optical stimulation for modulation of vasoconstriction in the deep brain of a living animal. Biomed. Opt. Express 8, 2781 (2017).

11. Skocek, O. et al. High-speed volumetric imaging of neuronal activity in freely moving rodents. Nat. Methods 15, 1–4 (2018).

12. Senarathna, J. et al. A miniature multi-contrast microscope for functional imaging in freely behaving animals. Nat. Commun. 10, 1–13 (2019).

13. Holloway, J., Wu, Y., Sharma, M. K., Cossairt, O. & Veeraraghavan, A. SAVI: Synthetic apertures for long-range, subdiffraction-limited visible imaging using Fourier ptychography. Sci. Adv. 3, e1602564 (2017).

14. Wu, Y., Sharma, M. K. & Veeraraghavan, A. WISH: wavefront imaging sensor with high resolution. Light Sci. Appl. 8, 44 (2019).

15. Boominathan, V., Adams, J., Robinson, J. & Veeraraghavan, A. PhlatCam: Designed phase-mask based thin lensless camera. IEEE Trans. Pattern Anal. Mach. Intell. 1–1 (2020) doi:10.1109/tpami.2020.2987489.

16. Betzig, E. et al. Imaging intracellular fluorescent proteins at nanometer resolution. Science 313, 1642–5 (2006).

17. Rust, M. J., Bates, M. & Zhuang, X. Sub-diffraction-limit imaging by stochastic optical reconstruction microscopy (STORM). Nat. Methods 3, 793–796 (2006).

18. Tian, L. & Waller, L. 3D intensity and phase imaging from light field measurements in an LED array microscope. Optica 2, 104 (2015).

19. Prevedel, R. et al. Simultaneous whole-animal 3D imaging of neuronal activity using light-field microscopy. Nat Methods 11, 727–730 (2014).

20. Ralston, T. S., Marks, D. L., Scott Carney, P. & Boppart, S. A. Interferometric synthetic aperture microscopy. Nat Phys 3, 129–134 (2007).

21. Ozcan, A. & Demirci, U. Ultra wide-field lens-free monitoring of cells on-chip. Lab. Chip 8, 98–106 (2008).

22. Adams, J. K. et al. Single-frame 3D fluorescence microscopy with ultraminiature lensless FlatScope. Sci. Adv. 3, e1701548 (2017).

23. Antipa, N. et al. DiffuserCam: lensless single-exposure 3D imaging. Optica 5, 1 (2018).

24. Greenbaum, A. et al. Imaging without lenses: achievements and remaining challenges of wide-field on-chip microscopy. Nat. Methods 9, 889–895 (2012).

25. Gill, P. R. & Stork, D. G. Lensless Ultra-Miniature Imagers Using Odd-Symmetry Spiral Phase Gratings. Imaging Appl. Opt. CW4C.3 (2013).

26. Asif, S., Ayremlou, A., Sankaranarayanan, A., Veeraraghavan, A. & Baraniuk, R. FlatCam: Thin, Lensless Cameras using Coded Aperture and Computation. IEEE Trans. Comput. Imaging 1 (2016).

27. Boominathan, V. et al. Lensless Imaging: A computational renaissance. IEEE Signal Process. Mag. 33, 23–35 (2016).

28. Horisaki, R., Ogura, Y., Aino, M. & Tanida, J. Single-shot phase imaging with a coded aperture. Opt Lett 39, 6466–6469 (2014).

29. DeWeert, M. J. & Farm, B. P. Lensless coded aperture imaging with separable doubly Toeplitz masks. Opt. Eng. 54, 023102 (2015).

30. Kuo, G., Antipa, N., Ng, R. & Waller, L. DiffuserCam: Diffuser-Based Lensless Cameras. in Imaging and Applied Optics 2017 (3D, AIO, COSI, IS, MATH, pcAOP) CTu3B.2 (2017). doi:10.1364/COSI.2017.CTu3B.2.

31. Sencan, I., Coskun, A. F., Sikora, U. & Ozcan, A. Spectral demultiplexing in holographic and fluorescent on-chip microscopy. Sci. Rep. 4, 3760 (2014).

32. Coskun, A. F., Sencan, I., Su, T.-W. & Ozcan, A. Lensless wide-field fluorescent imaging on a chip using compressive decoding of sparse objects. Opt. Express 18, 10510–23 (2010).

33. Pavani, S. R. P. et al. Three-dimensional, single-molecule fluorescence imaging beyond the diffraction limit by using a double-helix point spread function. Proc Natl Acad Sci U A 106, 2995–2999 (2009).

34. Chen, J., Hirsch, M., Heintzmann, R., Eberhardt, B. & Lensch, H. P. A. A Phase-coded Aperture Camera with Programmable Optics. Electron. Imaging 2017, 70–75 (2017).

35. Wu, Y., Boominathan, V., Chen, H. & Others. PhaseCam3D—Learning Phase Masks for Passive Single View Depth Estimation. 2019 IEEE (2019).

36. Chang, J., Sitzmann, V., Dun, X., Heidrich, W. & Wetzstein, G. Hybrid optical-electronic convolutional neural networks with optimized diffractive optics for image classification. Sci Rep 8, 12324 (2018).

37. Chi, W. & George, N. Optical imaging with phase-coded aperture. Opt Express 19, 4294 (2011).

38. Wang, W. et al. Generalized method to design phase masks for 3D super-resolution microscopy. Opt Express 27, 3799–3816 (2019).

39. Miyamoto, D. & Murayama, M. The fiber-optic imaging and manipulation of neural activity during animal behavior. Neurosci Res 103, 1–9 (2016).

40. Richard, C., Renaudin, A., Aimez, V. & Charette, P. G. An integrated hybrid interference and absorption filter for fluorescence detection in lab-on-a-chip devices. Lab. Chip 9, 1371–1376 (2009).

41. Badhiwala, K. N., Gonzales, D. L., Vercosa, D. G., Avants, B. W. & Robinson, J. T. Microfluidics for electrophysiology, imaging, and behavioral analysis of Hydra. Lab Chip 18, 2523–2539 (2018).

42. Dupre, C. & Yuste, R. Non-overlapping Neural Networks in Hydra vulgaris. Curr Biol 27, 1085–1097 (2017).

43. Juliano, C. E., Lin, H. & Steele, R. E. Generation of transgenic Hydra by embryo microinjection. J. Vis. Exp. JoVE 51888 (2014) doi:10.3791/51888.

44. Szymanski, J. R. & Yuste, R. Mapping the whole-body muscle activity of Hydra vulgaris. Curr Biol (2019).

45. Tzouanas, C. N., Kim, S., Badhiwala, K. N., Avants, B. W. & Robinson, J. T. Thermal stimulation temperature is encoded as a firing rate in a Hydra nerve ring. bioRxiv 787648 (2019) doi:10.1101/787648.

46. Dombeck, D. A., Graziano, M. S. & Tank, D. W. Functional Clustering of Neurons in Motor Cortex Determined by Cellular Resolution Imaging in Awake Behaving Mice. J. Neurosci. 29, 13751–13760 (2009).

47. Vanni, M. P., Chan, A. W., Balbi, M., Silasi, G. & Murphy, T. H. Mesoscale Mapping of Mouse Cortex Reveals Frequency-Dependent Cycling between Distinct Macroscale Functional Modules. J. Neurosci. 37, 7513–7533 (2017).

48. Barson, D. et al. Simultaneous mesoscopic and two-photon imaging of neuronal activity in cortical circuits. Nat. Methods 17, 107–113 (2020).

49. Pnevmatikakis, E. A. & Giovannucci, A. NoRMCorre: An online algorithm for piecewise rigid motion correction of calcium imaging data. J. Neurosci. Methods 291, 83–94 (2017).

50. Samaniego, A., Boominathan, V., Sabharwal, A. & Veeraraghavan, A. mobileVision. in Proceedings of the Wireless Health 2014 on National Institutes of Health - WH ’14 1–8 (ACM Press, 2014).

51. Candes, E. J., Li, X., Ma, Y. & Wright, J. Robust Principal Component Analysis? J. ACM 58, (2011).

52. Lin, Z., Ganesh, A., Wright, J. & Wu, L. Fast convex optimization algorithms for exact recovery of a corrupted low-rank matrix. Comput. Adv. Ldots 1–18 (2009) doi:10.1016/j.jsb.2012.10.010.

53. Bouchard, M. B. et al. Swept confocally-aligned planar excitation (SCAPE) microscopy for high-speed volumetric imaging of behaving organisms. Nat. Photonics 9, 113–119 (2015).

54. Ye, F., Avants, B. W., Veeraraghavan, A. & Robinson, J. T. Integrated light-sheet illumination using metallic slit microlenses. Opt. Express 26, 27326–27338 (2018).

55. Ghanbari, L. et al. Cortex-wide neural interfacing via transparent polymer skulls. Nat Commun 10, 1500 (2019).

56. Goldey, G. J. et al. Removable cranial windows for long-term imaging in awake mice. Nat Protoc 9, 2515–2538 (2014).

