## Supplementary Figures for "*In vivo* fluorescence imaging with a flat, lensless microscope"

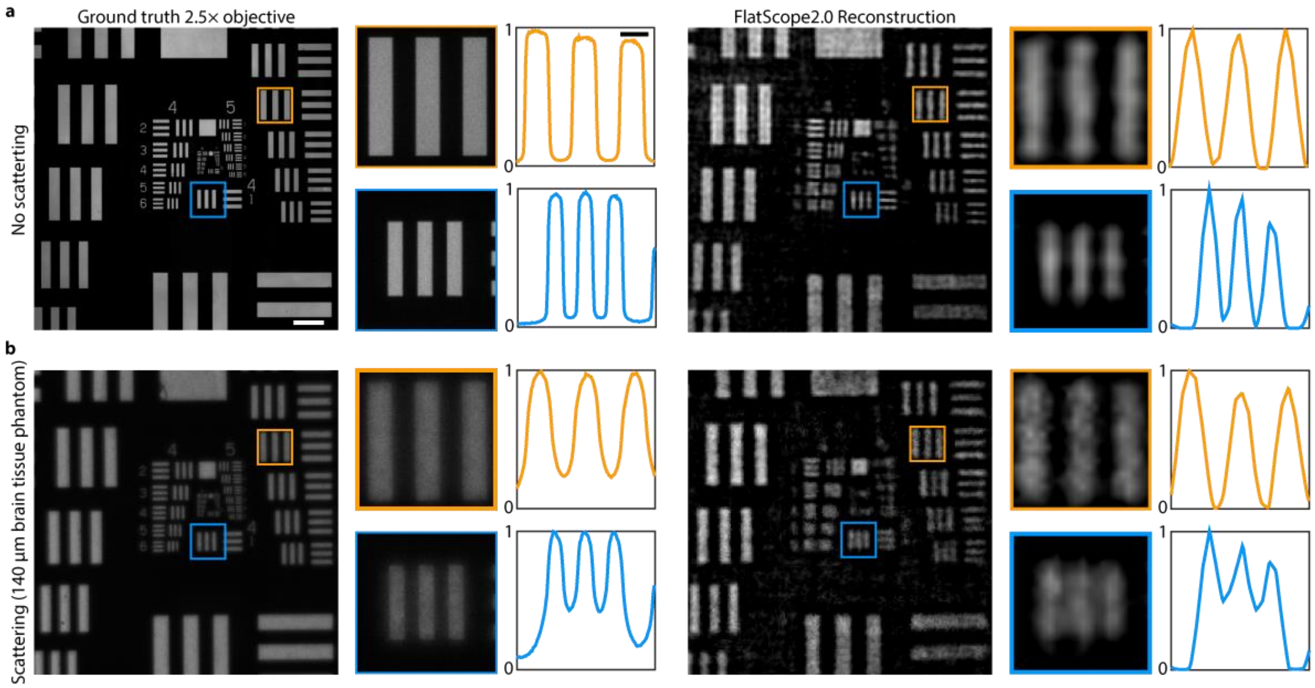


**Supplementary Figure 1 | Imaging through scattering medium. a**, Ground truth (left, captured with a 2.5× objective and FlatScope 2.0 reconstruction (right, captured at ~8.5 mm from the device) of USAF 1951 resolution target. Scale bar, 250 µm. Zoom-ins are shown for Group 3 element 3 and group 4 element 1. Scale bar, 50 µm. **b**, Ground truth (left, captured with a 2.5× objective) and FlatScope 2.0 reconstruction (right, captured at ~8.5 mm from the device) - scale bar, 250 µm) of USAF 1951 resolution target captured through 140 µm of brain tissue phantom. Scale bar, 250 µm. Zoom-ins are shown for Group 3 element 3 and group 4 element 1. Scale bar, 50 µm.

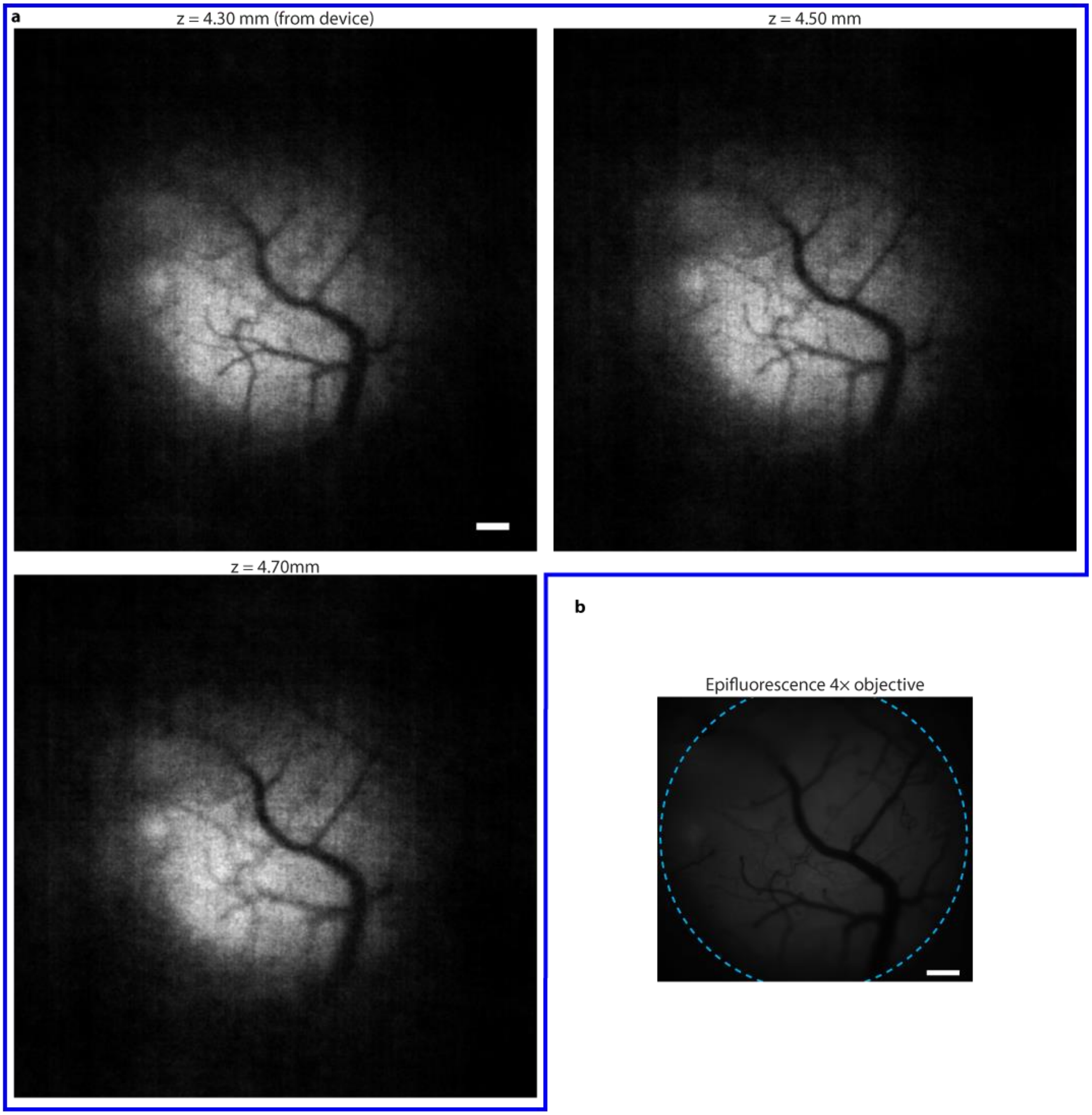


**Supplementary Figure 2 | Full FlatScope2.0 field of view for *in vivo* mouse brain recordings. a**, Full FOV FlatScope2.0 reconstructions of mouse brain in vivo at 4.3 mm, 4.5 mm, and 4.7 mm from the device. Dark areas (Around the center) are regions located outside of the cranial window. Scale bar, 250 μm **b**, Epifluorescence image of same region of the mouse brain captured with a 4× microscope objective. Dashed cyan line indicates the outer edge of the cranial window. Scale bar, 250 μm.

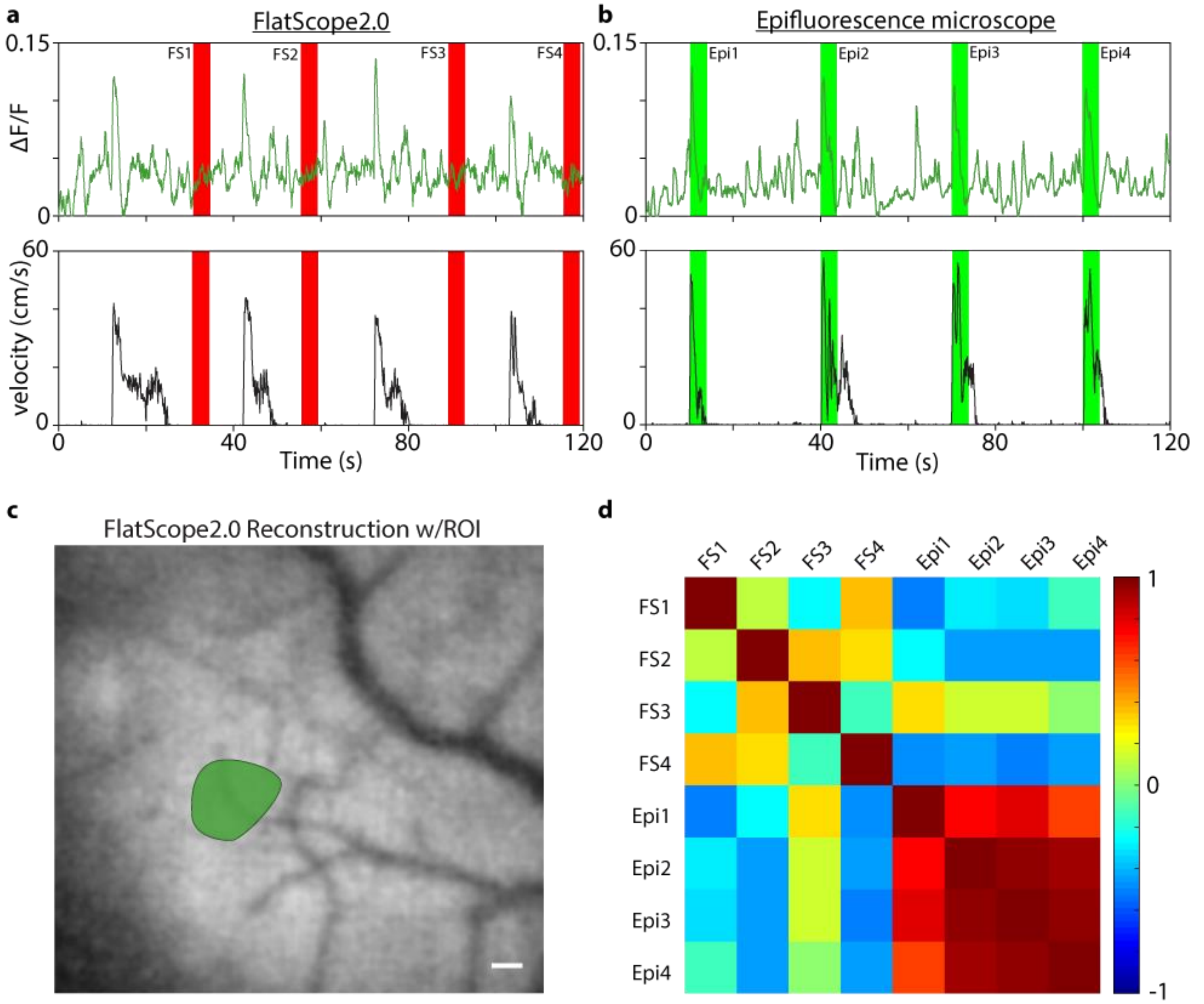


**Supplementary Figure 3 | Comparison of FlatScope2.0 $Ca^{2+}$ responses during stationary periods to stimulus-evoked responses in epifluorescence recordings.** **a**, ΔF/F trace and treadmill velocity for FlatScope2.0 during recording session. The 4-second windows are selected when the mouse is stationary (having a velocity < 1 cm/s), shown in red. **b**, ΔF/F trace and treadmill velocity for epifluorescence during recording session. The rising edge of the 4-second windows in green correspond to the application of tactile stimuli. **c**, FlatScope2.0 reconstruction with a single ROI of high-activity marked. Scale bar, 100 μm. **d**, Correlation matrix comparing the 4-second windows of little to no movement (red) and stimulus response activity (green). (FS - FlatScope2.0, Epi-Epifluorescence microscope).

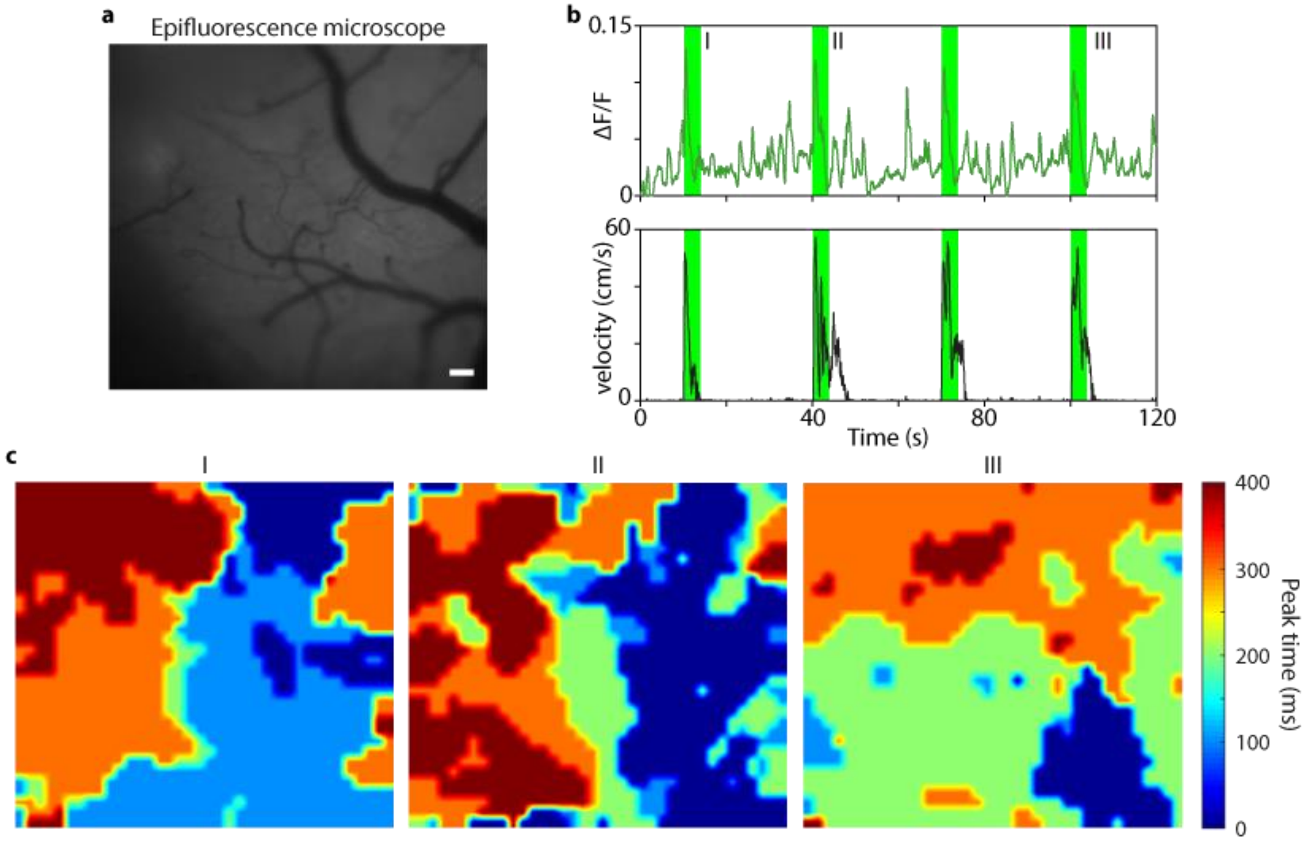


**Supplementary Figure 4 | Spatiotemporal $Ca^{2+}$ dynamics from epifluorescence recording. a**, Single frame from epifluorescence microscope capture of ROI. Scale bar, 100 μm. **b**, ΔF/F trace and treadmill velocity for epifluorescence during recording session. The rising edge of the 4-second windows in green correspond to the application of tactile stimuli. **c**, Heat maps for epifluorescence captures showing spatiotemporal $Ca^{2+}$ dynamics time-aligned with stimuli (at I, II, and III). Colormap shows the time at which pixels have their peak response for ΔF/F during a 400 ms period.

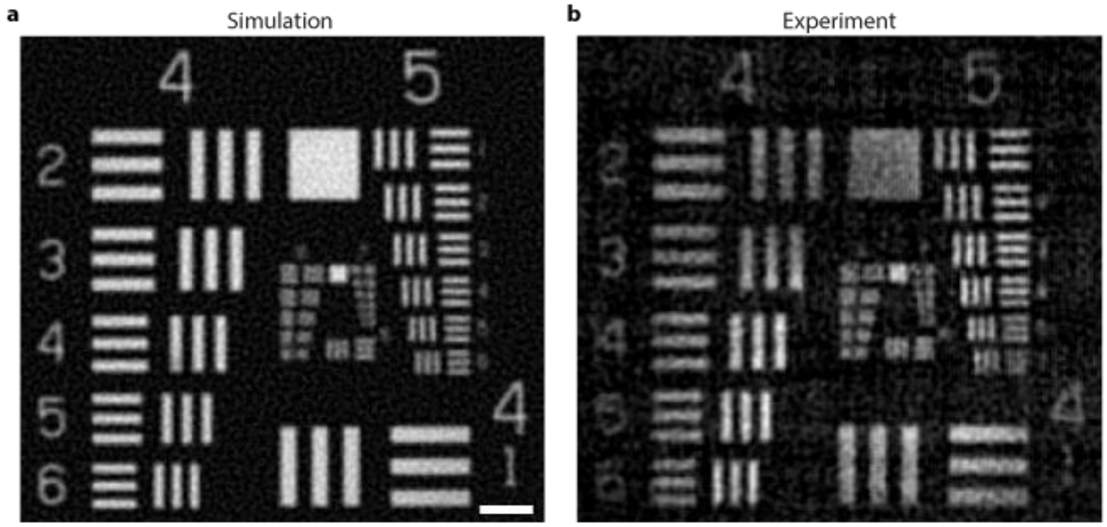


**Supplementary Figure 5 | Simulation vs experimental result of USAF. a,** Simulated FlatScope 2.0 reconstruction of USAF target at 4.15 mm from device. Scale bar, 100 μm. **b**, Experimental FlatScope 2.0 reconstruction of USAF target at the same distance from device. The simulation result shows a very close match to the experimental result with group 5 element 6 being resolved.

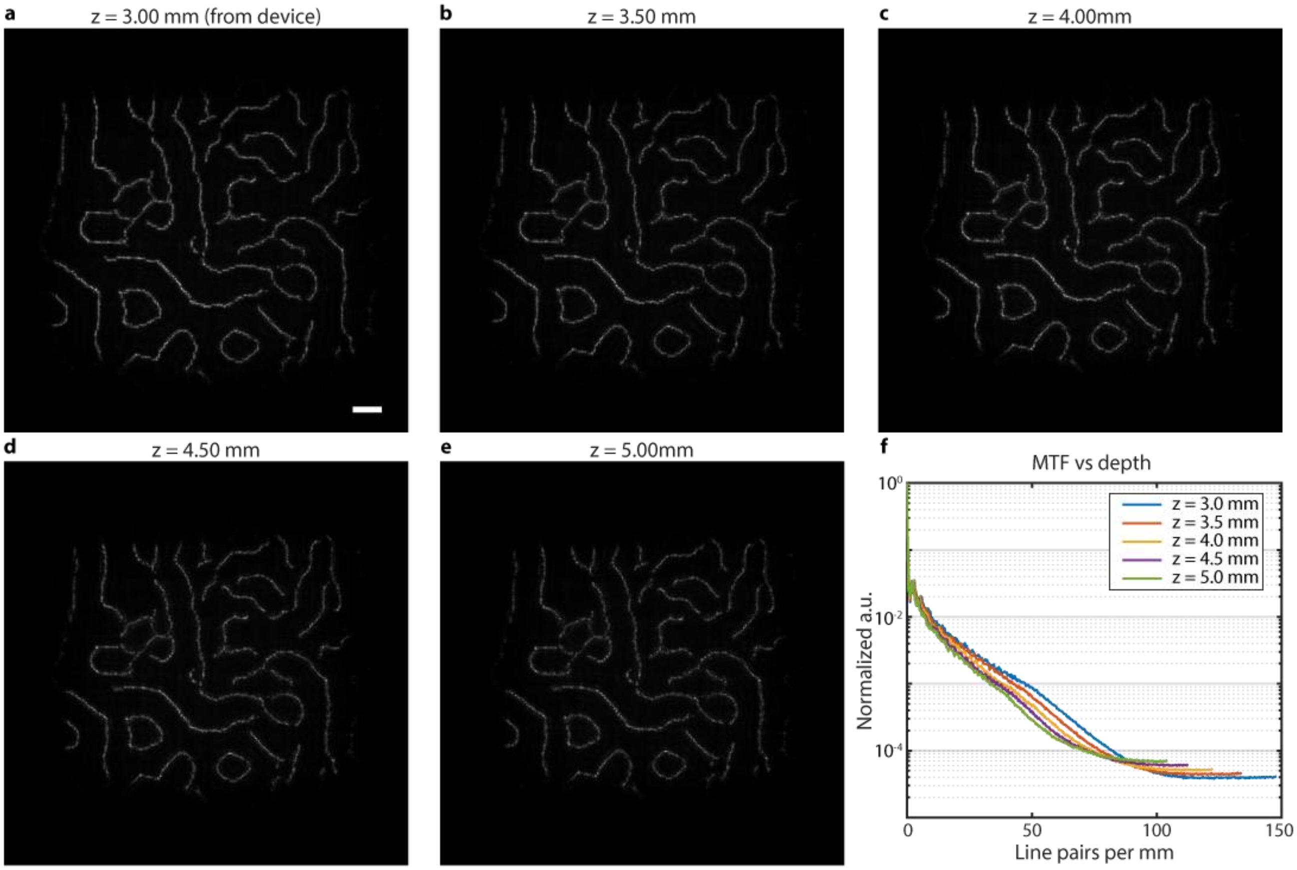


**Supplementary Figure 6 | Captured FlatScope2.0 point spread functions.** FlatScope 2.0 PSFs captured by imaging a 10 µm fluorescent microsphere (see Methods) at different depth planes of **a**, 3.00 mm, **b**, 3.50 mm, **c**, 4.00 mm, **d**, 4.50 mm, and **e**, 5.00 mm. Scale bar, 100 µm. **f**, Computed MTF from PSFs at different depths from FlatScope2.0. We observe that the resolution decreases with increasing depth, as seen from the shifting of MTF curves to the left.

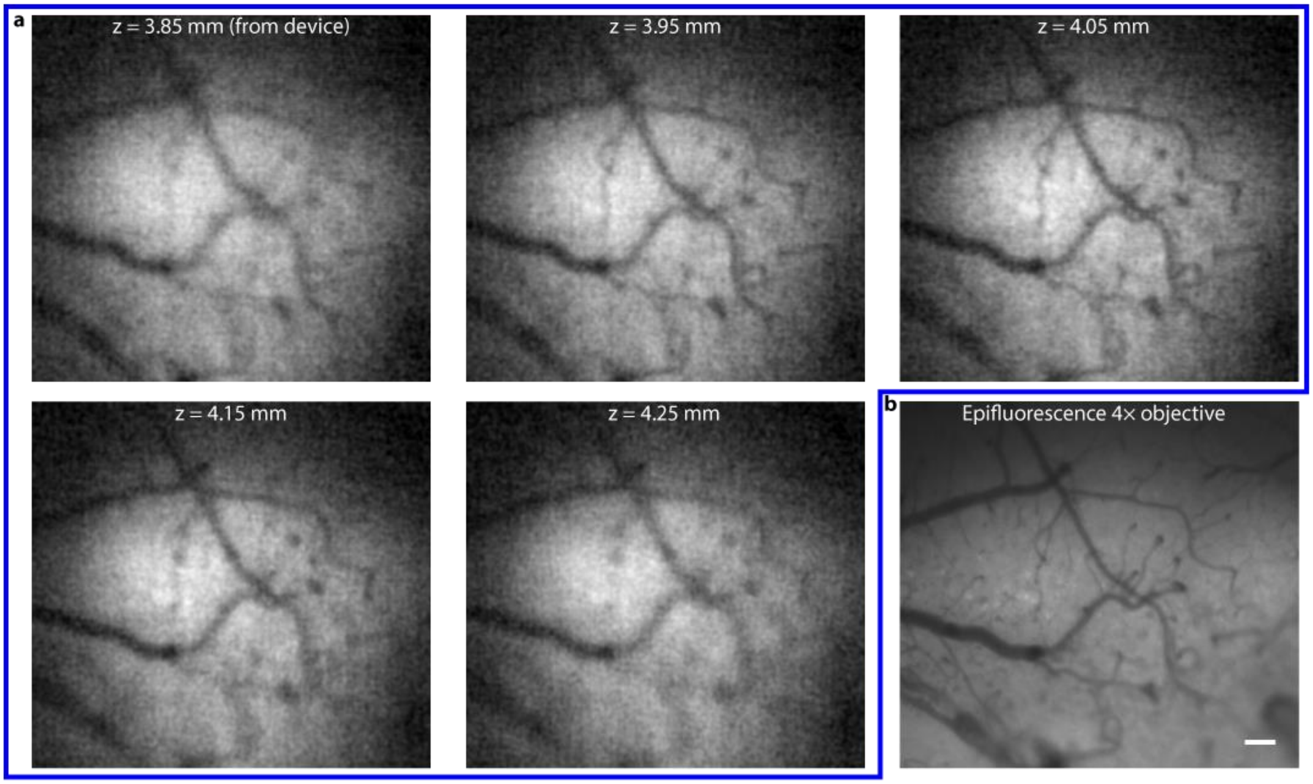


**Supplementary Figure 7 | Digitally refocusing in mouse brain. a**, FlatScope 2.0 reconstructions for five different depth planes. At 4.05 mm, the image becomes sharpest for the FOV. **b**, Epifluorescence image for the same FOV. Scale bar, 100 µm.

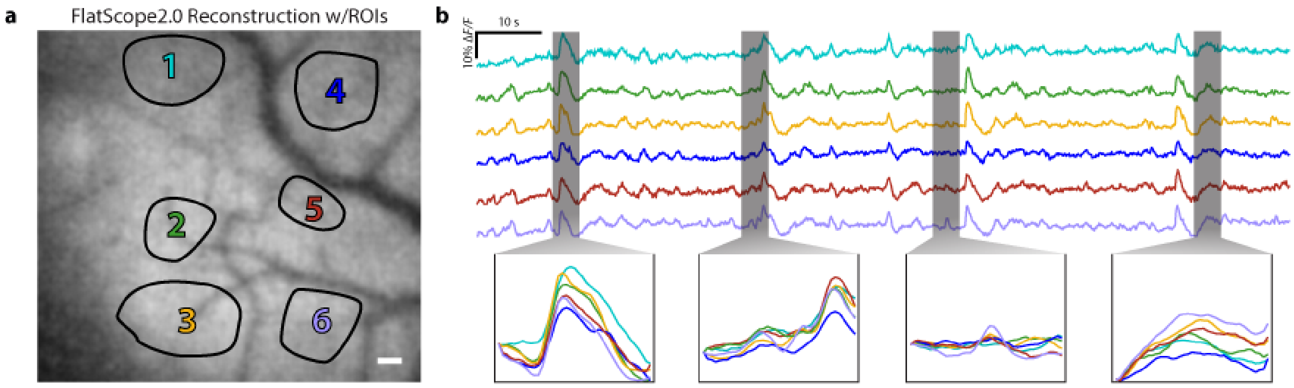


**Supplementary Figure 8 | $Ca^{2+}$ responses from Flatscope2.0 recording across multiple ROIs. a**, FlatScope2.0 reconstruction showing multiple ROIs. Scale bar 100 µm. **b**, ΔF/F traces for the multiple regions with zoom-ins showing differences in activity across ROIs.
